# Attention and prediction modulations in expected and unexpected visuospatial trajectories

**DOI:** 10.1101/2020.11.10.376368

**Authors:** Kristen S. Baker, Alan J. Pegna, Naohide Yamamoto, Patrick Johnston

## Abstract

Humans are constantly exposed to a rich tapestry of visual information in a potentially changing environment. To cope with the computational burden this engenders, our perceptual system must use prior context to simultaneously prioritise stimuli of importance and suppress irrelevant surroundings. This study investigated the influence of prediction and attention in visual perception by investigating event-related potentials (ERPs) often associated with these processes, N170 and N2pc for prediction and attention, respectively. A contextual trajectory paradigm was used which violated visual predictions and neglected to predetermine areas of spatial interest, to account for the potentially unpredictable nature of a real-life visual scene. Participants (*N*=36) viewed a visual display of cued and non-cued shapes rotating in a five-step predictable trajectory, with the fifth and final position of either the cued or non-cued shape occurring in a predictable or unpredictable spatial location. To investigate the predictive coding theory of attention we used factors of attention and prediction, whereby attention was manipulated as either cued or non-cued conditions, and prediction manipulated in either predictable or unpredictable conditions. Results showed both enhanced N170 and N2pc amplitudes to unpredictable compared to predictable stimuli. Stimulus cueing status also increased N170 amplitude, but this did not interact with stimulus predictability. The N2pc amplitude was not affected by stimulus cueing status. In accordance with previous research these results suggest the N170 is in part a visual prediction error response with respect to higher-level visual processes, and furthermore the N2pc may index attention reorientation. The results demonstrate prior context influences the sensitivity of the N170 and N2pc electrophysiological responses. These findings add further support to the role of N170 as a prediction error signal and suggest that the N2pc may reflect attentional reorientation in response to unpredicted stimulus locations.

## Introduction

Humans perceive their visual surroundings as a complete picture of the environment. In order for visual perception to take place, the brain must use prior information (prediction) to simultaneously guide focus to task relevant information whilst suppressing irrelevant information (attention). These two key characteristics of the visual environment are considered to influence successful visual perception. Attention involves enhancing task-relevant information in the environment, whilst simultaneously suppressing irrelevant information [1, 2]. In vision, the direction of attention refers to the prioritisation of visual objects, features or regions of space which are relevant to the viewers’ internal and external goals [3]. Prediction in cognition refers to the internal representations of stimulus likelihood based on prior experience [4]. Predictive coding models of perception stress the interplay of top-down and bottom-up signals in resolving percepts [4-6]. Such models suggest incoming stimulus input is constantly checked against predicted inputs; where a mismatch occurs, prediction error signals inform the updating of future predictions [5, 6]. In visual processing, this theory proposes an internal model of a scene is generated in higher order visual cortical areas, which receive feedback from actual visual input, turning the incorrect predictions into error signals. Subsequently, this leads to continuously updated predictions in a recursive hierarchical fashion [4, 6].

A prominent approach to investigate the cognitive mechanisms associated with prediction and attention in visual processing is to measure electrophysiological recordings time-locked to stimulus onset, termed event-related potentials (ERPs). Here we consider two widely studied visual ERPs – the N170 and the N2pc. The N170 has been of interest with respect to the processing of faces, and as the first ERP component indexing “higher-level” vision [4, 7]. Recent evidence has shown the N170 may be strongly modulated by violated expectations [8-11]. The N2pc, a lateralised difference wave potential occurring slightly later than the N170, has been traditionally associated with spatially selective attention, and is thought to index processes relating to prioritised processing of attended stimuli. Since recent work has shown the earlier occurring N170 is strongly modulated by predictions, here we address the question of whether the attention-relevant N2pc might show similar sensitivity [8, 9, 11].

### N2pc

The N2pc is a component considered to index attentional processes. The N2pc has been observed to salient lateralised targets in competition with distractors, in the cerebral hemisphere contralateral to the visual field of the target [12]. It is calculated as a contralateral-minus-ipsilateral difference waveform, with the resultant difference waveform indicative of the lateralised cognitive activity [13]. Initial evidence for the N2pc elicited to targets came from studies which used different combinations of simple visual objects (such as letters, shapes, and colours) with a target and distractor in opposing visual fields. Despite the presence or absence of distractors the N2pc was elicited contralateral to target stimuli (e.g., [14-16]). This suggested the N2pc is more dependent on target characteristics. Whilst the N2pc was initially investigated in basic visual processing [12, 14, 17], more recently the N2pc has been identified in studies of individual differences in attention as broad as rejection cues in low self-esteem [18], detection of threat in social anxiety [19, 20], face processing after exposure to violent video games [21], and fear related phobias [22].

Since the discovery of the N2pc a prominent view is this component reflects suppression of neighbouring distractors in favour of enhancing identification of the target. This is analogous to the spotlight metaphor of spatial attention where information in the beam of the spotlight receives enhanced processing [23]. The N2pc component is thought to index the operations of a spatial filter enhancing processing of a particular location in space whilst inhibiting the interference of surrounding distractors. This is supported by studies in which the N2pc was elicited to prospective cues of targets [24], repeated rather than novel arrays [25], and expected target locations marked by placeholder objects [26]. According to this idea, it would be expected the N2pc would be larger to a cued stimulus landing in a predictable region as a filter has pre-emptively been set up to accommodate expected processing in this location.

A contrasting view considers the N2pc to reflect the reorientation of attention. In one study [27], two targets were shown in quick succession with the second target (T2) appearing in the same or opposite visual field of the first target (T1). An attenuated N2pc was found to T2 when it followed the same visual field as T1, however when T2 was in the opposite field a second N2pc occurred. The N2pc has also recently been found to be elicited to orientation singletons which swap as target and distractor between trials [28]. This evidence suggests the N2pc is reflective of the reorientation of attention rather than an enhancement of sustained target processing in a predetermined spatial location.

### N170

The N170 is a negative deflection of the ERP response observed post stimulus onset at approximately 150 to 200 milliseconds (ms) in the lateral occipital region [29]. Early research examining the characteristics of the N170 focussed on its role in indexing stimulus category, demonstrating an enhanced sensitivity to faces including configuration and emotional expression [30-33]. Other studies suggest that the N170 is not exclusively face-specific [34], with recent evidence using face stimuli in addition to shapes, bodies, and nude statuettes demonstrating the N170 could be an index of mismatches between prior expectations and actual stimulus input in higher-level vision [9, 11]. Johnston and colleagues [9] demonstrated this in experiments in which they reported a novel contextual trajectory paradigm, whereby participants observed sequences of images in either predictable or unpredictable conditions, with the final step of unpredictable conditions violating the established order of the sequence of preceding stimuli. Investigating both EEG and magnetoencephalography (MEG) responses, the N170 and M170 were larger in amplitude for unpredictable versus predictable stimulus onsets, irrespective of object category. Careful control of stimulus presentation ensured that these effects must be due to contextually based expectations rather than final stimulus transitions themselves, as these were identical across conditions. Furthermore, N170 amplitudes to unexpected stimuli have been found to be enhanced in a dose dependent manner, based on the strength of accumulated evidence for the predicted outcome, and the degree to which the observed stimulus differs from that outcome [11]. These studies support the idea the N170 is, at least in part, a visual prediction error signal reflecting a conflict between internally generated predictions and actual sensory input [9, 11]. In light of the recent findings of the N170 modulated by prediction error processes, and given the close proximity in electrode sites (N170 often P7 and P8, N2pc often PO7 and PO8), and time windows (N170 peaks at 170ms, N2pc generally varies between 180-300ms) [29, 33] of the components, the question is raised, of whether the N2pc, like the N170, might be modulated by stimulus predictability.

### The present study

The present study was designed to determine the influence of expectancy violations of prediction and attention in visual processing by analysing responses of the N170 and N2pc. To accurately measure discrete early visual temporal processes, our paradigm combined predictable and unpredictable visuospatial trajectories. Participants viewed a five-step sequential trajectory of a cued shape and non-cued shape, which concluded in a predictable or unpredictable spatial location. This allowed us to attempt to dissociate processes indexed by the N170 and the N2pc as markers of predictive spatial filtering or attention reorientation.

Based on previous findings, it was expected enhanced N170 amplitudes would exist for unpredictable compared to predictable conditions – however, there have been no previous studies examining whether or not this effect is modulated by attentional cueing. In relation to the N2pc, to date, there have been no studies employing a contextual trajectory method that induces (and violates) expectations regarding stimulus location. Therefore, it is not clear whether the N2pc simply reflects the actions of an attentional spatial filter, or whether it might also be modulated by expectancy violations. In accordance with the view that attention increases the precision of prediction error signals, it was hypothesised we would observe enhanced ERP amplitudes to cued versus non-cued and unpredictable compared to predictable stimuli, with the largest amplitude to unpredictable attended conditions [35-39]. On the other hand, if a particular ERP component indexes either only locus of attention or a mismatch between the observed and expected stimulus, then the ERP component should only be modulated by one and not the other of these factors [40-42].

## Method

### Participants

Data were initially collected from 40 participants; however, four participants were excluded from further analyses due to artefacts in the overall recording or in the electrodes of interest. Thus, 36 participants remained in the sample aged from 17-60 years (*M* = 25.25, *SD* = 9.29, 26 females, 29 right-handed). As there is minimal research of the N2pc in relation to the predictive coding theory, this sample size was based on previous studies that investigated predictability effects on the N170 and N2pc [11, 27, 43]. Participants received a gift card or university course credits for participation. All participants had normal or corrected-to-normal vision, and no history of neurological conditions. Ethical approval was granted by Queensland University of Technology’s Research Ethics Committee, and participants gave informed written consent.

### Stimuli and paradigm

For each trial participants viewed a visual display consisting of three black shapes (circle, square, and arrow), on a HP monitor with screen resolution of 1920×1080 pixels. The visual display consisted of an arrow positioned in the centre, pointing at one of the simultaneously presented shapes. The stimuli were arranged such that the circle and square were displayed in one of 12 different possible positions in a circular array. The arrow was always centrally fixed but varied in orientation in 30-degree increments, dependent on the shape it was pointing to. The arrow stimuli subtended approximately 1.9° of visual angle, with circle and square stimuli subtending 3°, offset by approximately 1.8° from the central fixation point. The square stimuli measured 225 × 225 pixels, the circle diameter 225 pixels, and the arrow stimuli 50 (length) x 24 (maximum width) pixels. Each sequence was made up of five trials whereby the first trial began with the cued shape and non-cued shape appearing at any of the 12 possible positions, followed by four additional steps in a clockwise or anti-clockwise trajectory circling around the arrow. The orientation of the arrow varied such that it always pointed at the position of the cued shape for each of the five steps in the sequence, except for the fifth step in unpredictable conditions whereby the arrow continued in the expected trajectory and the cued shape or non-cued shape moved back one position. Arrows cues have been found to automatically orient attention, even when being uninformative or incongruent to the correct location of cues [44, 45]. Hence, in each sequence the shape the arrow was pointing to was considered the cued shape, whereas the non-cued shape was the opposing shape the arrow was not pointing to. When the circle was the cued shape, the square was the non-cued shape and vice versa. As such there were equal presentations of circles and squares as both cued and non-cued shapes. Visual stimuli at the final step were situated with the cued shapes in the vertical midline and the non-cued shapes laterally, or vice versa (cued lateral, non-cued midline).

A duplicate set of images with red dots placed in the centre of the arrow were produced for red dot trials, which were to ensure participants’ gaze was maintained centrally throughout the duration of the task [8-11, 46]. Participants were asked to press the spacebar as soon as the red dot appeared. These trials appeared randomly in 10% of all five-trial sequences. The red dots were sized 12 pixels in diameter. Participants were required to respond correctly to more than 90% of red dot trials as indication of sufficient attention to the task [9]. Red dot trials were separately coded and then subsequently discarded during EEG pre-processing. Refer to S1 Fig for one example of the red dot trials.

For this study we decided upon four different conditions, in order to manipulate prediction and attention in accordance with previous similar studies whereby attention is often manipulated as attended and unattended conditions, and prediction manipulated as expected or unexpected conditions [35, 38, 40, 41]. Prediction was manipulated in either a predictable or unpredictable fashion, and attention was manipulated via cued and non-cued conditions. As such, there were four conditions termed predictable cued, unpredictable cued, predictable non-cued, and unpredictable non-cued. These were named with respect to whether the final location of either the cued or non-cued shape in the sequence was in a lateralised predicted or unpredicted position. Each condition was equally divided on rotation direction (clockwise or anti-clockwise), lateralised endpoint shape status (cued or non-cued), and endpoint position (left or right visual field). The four conditions were equated in low-level visual properties of the two shapes in the critical final trial, such that any differences in ERPs could be attributed to the preceding steps. There were 480 five-trial sequences in total, consisting of 108 sequences per condition and 48 red dot sequences. All conditions were presented randomly per participant. Each trial in the sequence was presented for 500ms, 0ms intertrial interval, and a 500ms intersequence interval. Refer to Fig 1 for a visual depiction of each of the conditions.

**Fig 1.**
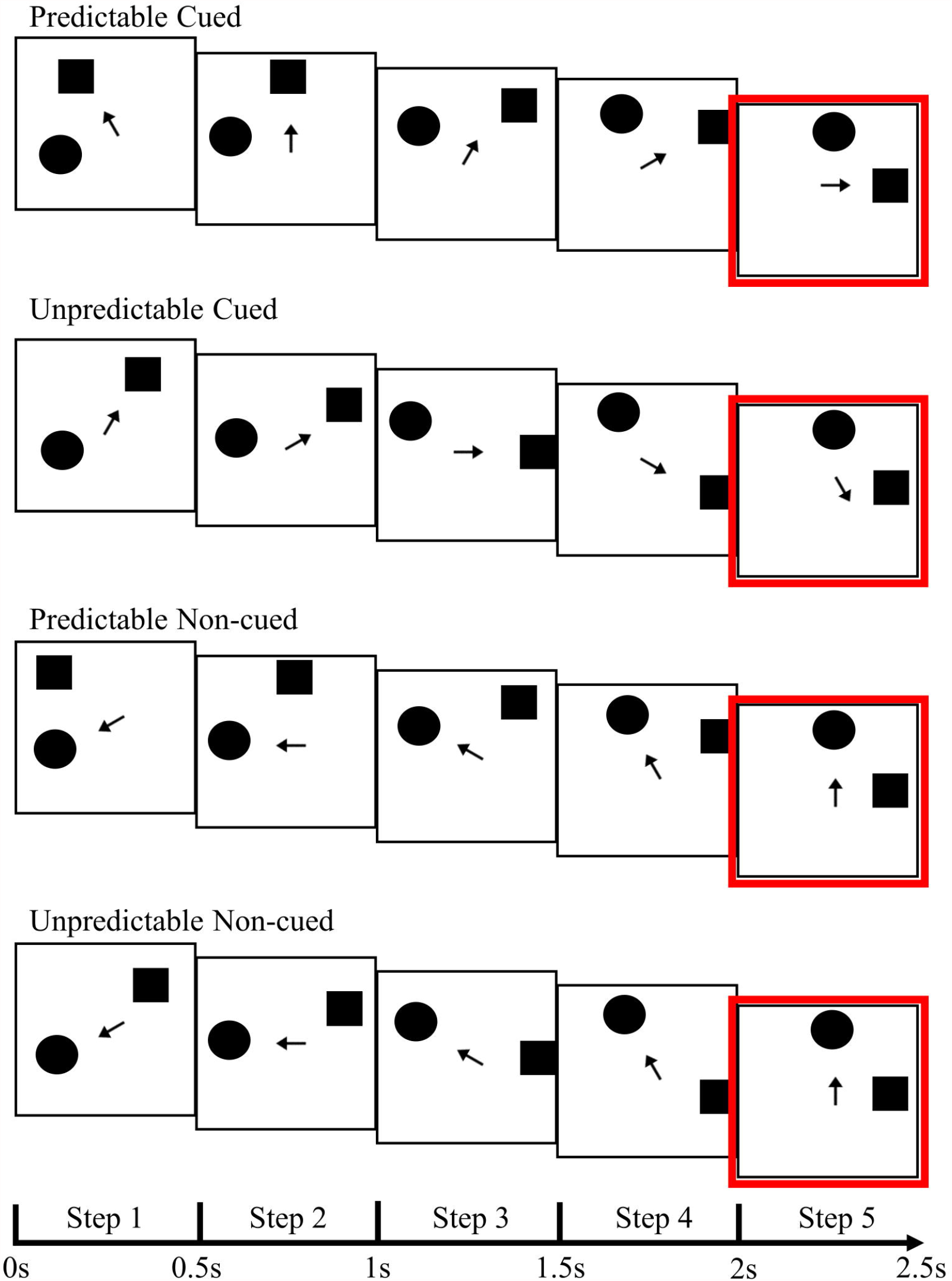
Examples of all conditions. Example of one sequence for each of the four conditions. Stimulus duration as indicated, 0ms intertrial interval, 500ms intersequence interval. Conditions are named with respect to lateralised shape status in the fifth trial, as outlined in red. All conditions were equal in low-level visual properties of the two shapes (excluding the arrow) in the critical final trial.

#### Procedure

Participants were seated approximately 60cm in front of the computer screen, fitted with the EEG cap and instructed to watch the sequence of images appearing on the screen. EEG data were recorded from 64 channels, using the international 10-20 system for electrode placement. Participants viewed a display of 480 sequences with each sequence consisting of the presentation of five shape trials. Participants viewed four blocks of 120 sequences each, with a break in between each block whereby they initiated the next block via button press. To ensure gaze was fixated centrally, participants were informed to fixate on the centre of the computer screen and respond via button press if a red dot appeared in the centre of the arrow. Otherwise, no behavioural response was required while viewing the trials.

### Data analysis

A BioSemi 64 channel amplifier recorded electrophysiological data at a sampling rate of 1024 Hz, using a common reference (CMS/DRL). EEG data was displayed to the experimenter on the BioSemi acquisition program, Actiview [47]. Electrode offset was maintained below 20 mV. Pre-processing of EEG data was performed with the software BrainVision Analyzer 2 [48]. Following this, ERP waveform processing was undertaken with MATLAB [49] and descriptive/inferential statistics with SPSS software [50].

#### Pre-processing

A band-pass filter between 0.1 to 30 Hz (24db/octave slope) with a notch filter of 50Hz was applied. Artefact rejection was performed by applying criterion of Min-Max fluctuations of >500µV in 100ms intervals, marking 100ms before and 100ms after the event. Visual inspection of the raw EEG data was then carried out and noisy electrodes were interpolated using the spherical spline interpolation method. Due to excessive noise and EEG recording artefacts, the aforementioned four participants were excluded at this stage with 36 remaining participants. Eye-blinks were identified and rejected with BrainVision Analyzer’s automated ocular correction procedure using independent components analysis. Data were re-referenced to an average electrode. Next, data were segmented into 2600ms time-locked epochs, beginning 100ms pre the first stimulus onset in the sequence and ending 500ms post final stimulus onset. Separate epochs were created for each condition as to whether the final stimuli were predictable or unpredictable, cued or non-cued, and lateralised shape ending in the left or right visual field. Next, an average of each epoch was generated. Due to the absence of an interstimulus interval final averaged waveforms were baseline corrected to the time period of stimulus onset of −150ms to 0ms of the average of both the fourth and fifth step in the sequence [9]. Grand average ERP waveforms were then generated from activity at electrode clusters for each of the four conditions, also divided by visual field, resulting in eight separate grand averages. Electrode clusters consisted of channels P5, P7, PO3, and PO7 for the left hemisphere (LH), and electrodes P6, P8, PO4, and PO8 for the right hemisphere (RH). These electrodes were chosen based on a review of typical sites of maximal N170 and N2pc responses in previous literature [9, 20, 29, 51, 52]. N170 responses were measured by pooling activity for both visual fields irrespective of the location of the final stimuli, for each condition. N2pc components were generated by subtracting activity in the electrode cluster of the corresponding ipsilateral hemisphere to stimulus presentation from activity in the contralateral hemisphere, resulting in a difference waveform.

## Results

Participants were included if they correctly identified >90% of red dot sequences. No participants were excluded on this basis, thus 36 participants remained. The remaining 36 participants achieved an accuracy rate of 98.43% correctly identifying red dot trials, with a mean reaction time of 497.64ms (*SD* = 12.97ms). EEG data were comprised of ERP waveforms at the fifth and final trial in each sequence, for each of the four conditions. The N170 peak-to-peak amplitude was measured by subtracting the N170 inflection points maxima from the P1 inflection points minima, as per previous studies [8, 11]. N170 amplitudes were measured in the time window 140ms to 210ms, and the P1 80ms to 130ms. The N2pc component was measured as the average amplitude between 200ms to 300ms post final stimulus onsets, as the difference wave (contralateral electrode cluster minus ipsilateral electrode cluster). All amplitudes reported herein were measured in microvolts (µV).

To investigate the hypotheses two omnibus repeated-measures ANOVAs, one for N170 and another for N2pc, were performed, each with factors of expectation (predictable and unpredictable) and shape status (cued and non-cued). Cook’s distance was calculated to identify potential outliers, with no outliers identified. The assumption of sphericity was met for both ANOVAs.

### N170

There was a significant main effect of expectation, *F*(1,35) = 8.62, *p* = .006, η_p_^2^ = .198. The N170 amplitude was significantly larger (i.e., more negative) to unpredictable trials (*M* = −2.29, *SE* = .24) compared to predictable trials (*M* = −2.02, *SE* = .20). There was a significant main effect of shape status, *F*(1,35) = 7.70, *p* = .009, η_p_^2^ = .180, with cued trials eliciting a larger N170 amplitude (*M* = −2.26, *SE* = .23) compared to non-cued trials (*M* = −2.05, *SE* = .21). The interaction effect was not significant, *F*(1,35) = 0.37, *p* = .548, η_p_^2^ = .010. These results indicate a larger N170 was elicited by lateralised cued shapes than lateralised non-cued shapes, and by unpredictable trials than predictable trials, however the effect of (un)predictability on N170 was not greater when it was a cued than non-cued shape. Fig 2 illustrates the grand averaged N170 scalp topographies and grand averaged waveform amplitudes.

**Fig 2.**
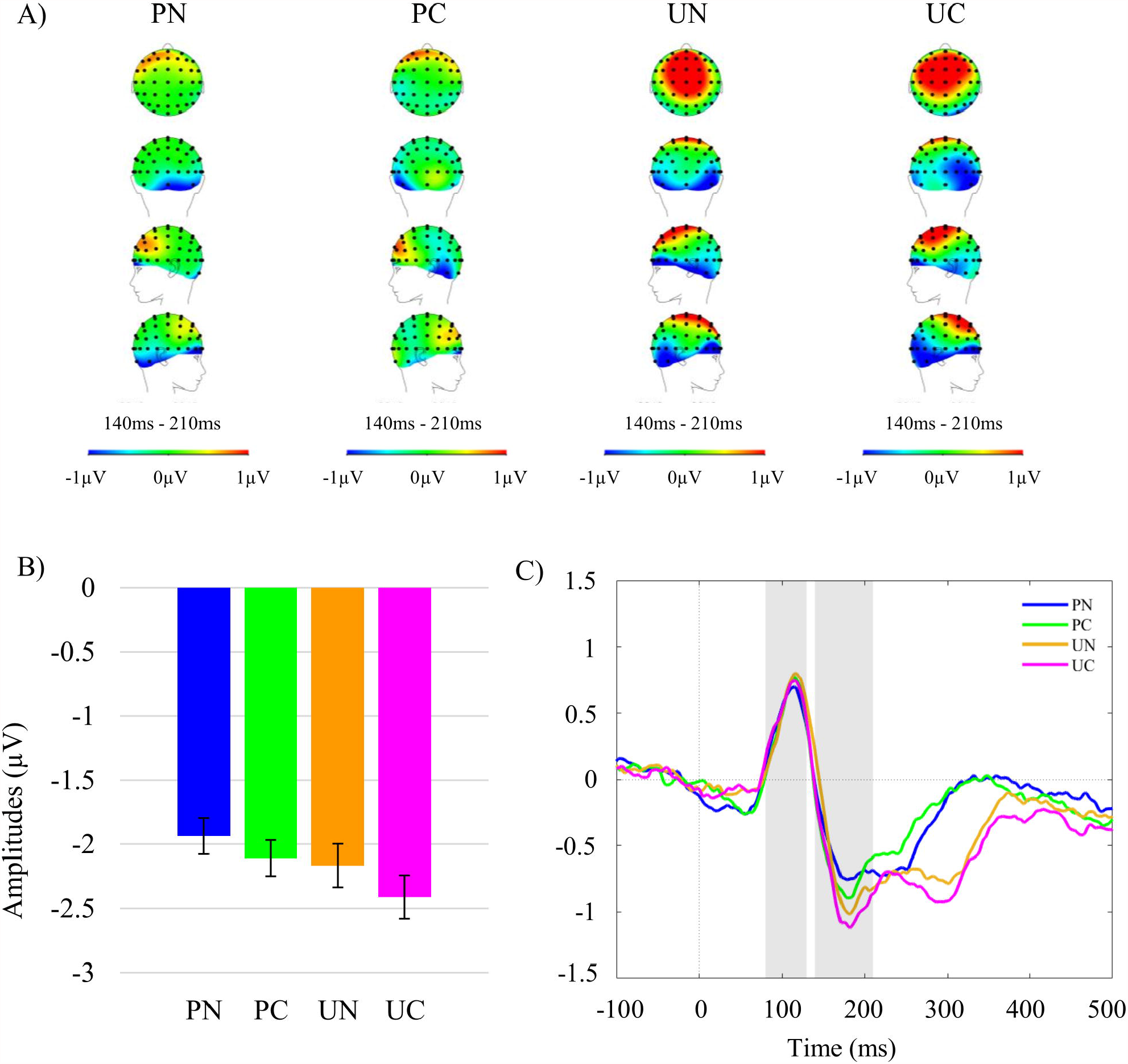
Grand average N170 topographies and amplitudes. A) Scalp topographies across all electrodes during the N170 time window for all conditions. From top to bottom: top, back, left, and right view. Heat map scales were set at −1µV minimum and 1µV maximum for display purposes. B) Grand averaged mean N170 amplitudes (µV) for all conditions pooled across left and right hemisphere electrode clusters. Error-bars denote 95% confidence intervals calculated for repeated measures using the Cousineau-Morey Method [53]. C) Grand averaged waveforms pooled across hemispheres for all four conditions. Grey panels indicate the time windows tested with P1 80-130ms, and N170 140-210ms. PN= predictable non-cued, PC= predictable cued, UN= unpredictable non-cued, and UC= unpredictable cued trials.

### N2pc

There was a significant main effect of expectation, *F*(1, 35) = 4.98, *p* = .032, η_p_^2^ = .125. The N2pc amplitude was significantly larger (i.e., more negative) to unpredictable trials (*M* = −.10, *SE* = .08) compared to predictable trials (*M* = .19, *SE* = .12). There was no significant main effect of shape status, *F*(1, 35) = 0.023, *p* = .881, η_p_^2^ = .001, and no significant interaction effect *F*(1, 35) = 1.48, *p* = .232, η_p_^2^ = .041. These results demonstrate larger N2pc components occur to unpredictable conditions, but this is not contingent upon cued or non-cued status. Refer to Fig 3 for visual depictions of grand averaged ERP difference waveforms of the N2pc for all conditions.

**Fig 3.**
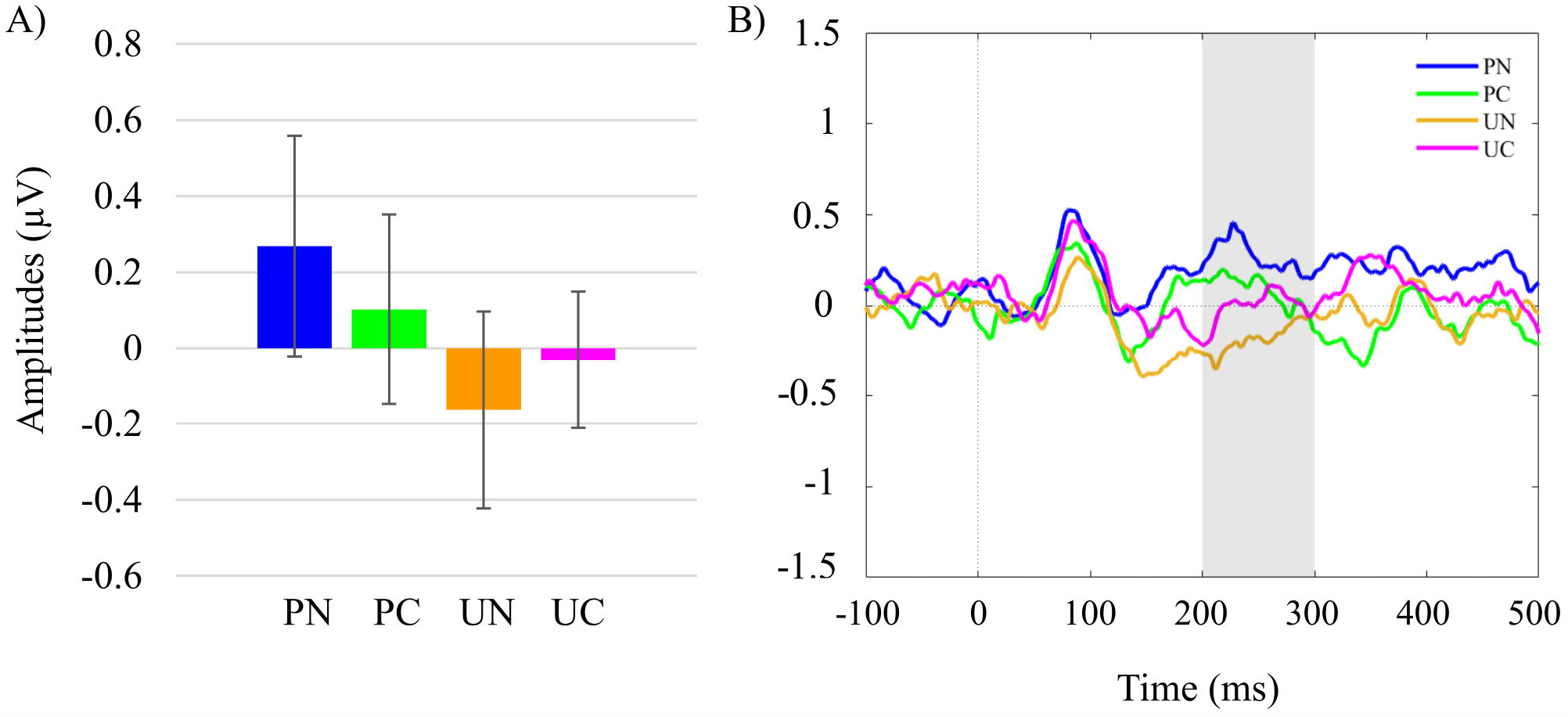
Grand average N2pc amplitudes. A) Grand averaged N2pc amplitudes (µV) pooled across hemispheres for all conditions (pooled contralateral activity minus pooled ipsilateral activity). B) Grand averaged ERP amplitudes (µV) of N2pc difference waves for all four conditions. Error-bars denote 95% confidence intervals calculated for repeated measures using the Cousineau-Morey Method [53]. Grey panels indicate the time windows tested of the N2pc component from 200-300ms. PN= predictable non-cued, PC= predictable cued, UN= unpredictable non-cued, and UC= unpredictable cued trials.

### Supplementary Analysis

The main analysis was informed by literature with respect to the conventional measurement of the N2pc in which a particular time window is often a priori specified as the focus of interest ranging from 180-200ms to 300ms [12, 13, 54-57]. To explore the robustness of these findings we conducted supplementary analyses adopting a temporal cluster permutation method [10, 58]. We constrained ourselves to examining subtraction waveforms across the pooled channels which were the focus of the main analyses and performed temporal cluster t-tests comparing (respectively) predictable versus unpredictable trials and cued versus non-cued trials. For each of these comparisons, we first performed t-tests at each time point between −100 to 500ms following stimulus onset. We set a cluster-forming height threshold of *t*(35) = +/−2.02 (equivalent to an uncorrected two-tailed alpha threshold of *p* <. 05) and treated consecutive time points exceeding this threshold as forming temporal clusters. Cluster values were generated by summing the t-values associated with each timepoint within the cluster. These cluster values were tested for significance against a null distribution of pseudo-clusters that were formed using a sign-flip permutation method to the subtraction waveforms of unpredictable minus predictable conditions, and separately cued minus non-cued conditions. For each of 10,000 permutations, each subtraction waveform was sign-flipped (positive values become negative, and vice versa) with a probability of *p* = .5. A t-test was then performed for each time-point, comparing against an expected value of zero. Temporal clusters were defined in the same way as previously described, and the same manner of calculating cluster-values was applied. For each permutation, the absolute value of the largest (positive or negatively valued) cluster value was added to the null distribution (this maximal pseudo-cluster-value technique implicitly controls for multiple comparisons [59]; also note that taking the absolute maximal value (positive or negative) makes the selected alpha criterion two-tailed). We set the alpha criterion as *p* <. 05 (two-tailed, corrected for multiple comparisons), thus, any cluster value exceeding the value of the 95^th^ percentile of the null distribution was considered significant. For shape status (cued versus non-cued trials) there were no clusters of time-points surviving the initial height threshold. For predictable versus unpredictable conditions there was a temporal cluster between 173ms-239ms following stimulus presentation with a cluster value of summed-t = 183.22. The 95^th^ percentile of the null distribution was summed-t = 30.34, thus, this value exceeds the *p* <. 05 threshold for significance. In fact, the estimated likelihood of a cluster of this magnitude under the null hypothesis was *p* < .00013. Refer to Fig 4 for depictions of the grand averaged waveforms and t statistics of predictable compared to unpredictable trials.

**Fig 4.**
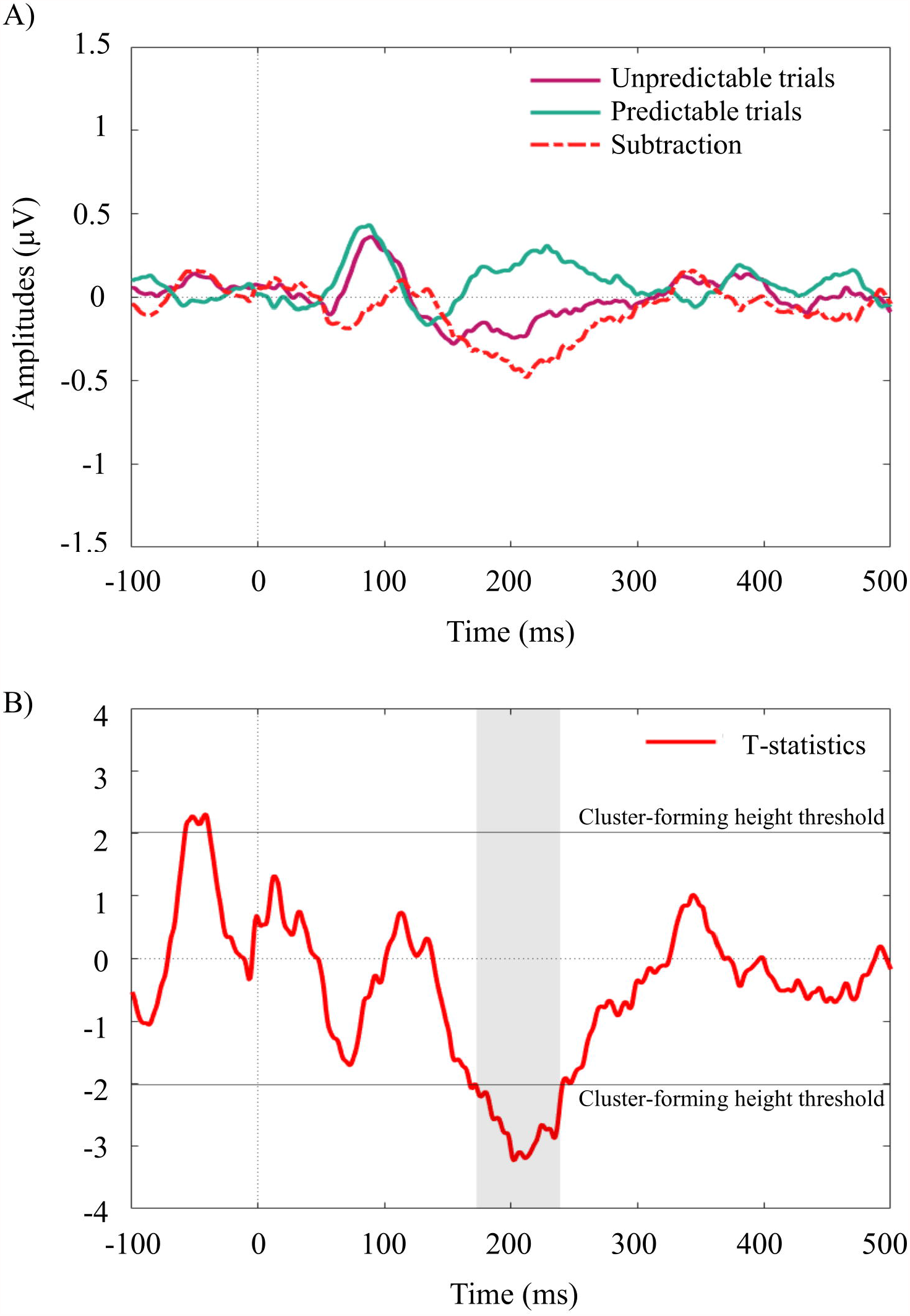
Temporal cluster permutation. A) Grand averaged N2pc amplitudes (µV) pooled across both predictable conditions and both unpredictable conditions, and the subtraction (difference) waveform of unpredictable minus predictable conditions. B) T statistics for the predictable and unpredictable conditions analysed with the temporal clustering permutation method. The sole significant temporal cluster from 173ms to 239ms is denoted by grey shading.

## Discussion

The aim of the present study was to investigate modulations of prediction and attention in visual processing using expectancy violations by analysing responses of the N170 and N2pc. We investigated the relationship between non-cued and cued stimuli using ERPs considered responsible for indexing these cognitive processes. It was shown that the N170 was modulated both by stimulus predictability (with larger amplitudes to unexpected stimulus locations) and by stimulus cueing (showing a greater amplitude to cued stimuli). There was no evidence to support an interactive effect of these factors. With respect to the N2pc, it was demonstrated that stimulus predictability modulated the ERP amplitude (again, with larger amplitude responses to unexpected stimulus locations), however, there were no effects of stimulus cueing, and no interactions.

The result of the N170 being larger in unpredictable conditions compared to predictable conditions support the view the N170 is in part reflective of a prediction error signal [9]. This study also supports the notion the N170 is attention dependent too, with the larger N170 found to cued stimuli and the smaller to non-cued stimuli as noted in previous research with N170 amplitudes modulated by cued stimuli [60-62]. This observation is not entirely unexpected since previous work [63] has shown both P1 and N1 ERPs to be enhanced by spatial attention cueing. Interestingly though, the two factors of stimulus predictability and stimulus cueing did not interact. This indicates, at this stage of processing at least, that the two processes may be independent, and even subserved by different neural substrates.

For the N2pc, the component was larger to unpredictable conditions compared to their predictable counterparts. Indeed, for predictable conditions it is not clear there are N2pc components, since the group means for the contralateral minus ipsilateral waveforms do not show a negativity during the N2pc time window. We interpret this pattern of results to indicate that the N2pc indexes processes relating to the reorientation of attention. Since there is no discernible N2pc to stimuli in predictable locations, this rules out the possibility that the N2pc is related to the pre-allocation of attention to specific spatial locations. Rather, the fact it is largest to unpredictable stimulus locations favours the interpretation that attention is redirected. These findings are consistent with other studies which found attenuation of the N2pc in a primed location [27], and reorientation to stimuli of alternating salience status between target and distractor [28]. The significant main effect for predictability demonstrates prediction plays an important role in the elicitation of the N2pc, which should be taken into consideration in future studies. This reiterates the importance of controlling for prior context and priming existent in paradigms which could influence the electrophysiological activity in cued and non-cued stimulus recognition. Given the observation of the N2pc to be modulated by attention redirection, future research should investigate how the amplitude of the N2pc component could be sensitive to manipulations of the dosage of violation, as previously found to vary for the N170 component [11].

That there was no significant effect of shape status (cued versus non-cued) on the N2pc component suggests the stimuli were not differentiated by attentional cued relevance. However, there was a main effect of shape status with the N170 which demonstrates differences were detected between cued and non-cued stimuli. This finding of a main effect of the N170 suggests the cueing arrow stimuli in this paradigm were effectively manipulating endogenous attentional relevance. Whilst predictability modulated N2pc amplitudes, but shape status (implicit cued or non-cued) did not, this suggests the N2pc in this paradigm has been modulated by redirection of attention irrespective of shape status. One possible explanation for this considers previous research whereby stimuli evoked an N2pc when specific stimuli shapes swapped between being targets and distractors on different trials [28]. Our results raise the question of whether previous findings may have been driven more by unexpected changes in attributes of the target stimulus rather than simply the status of being classified as a “target”.

The prediction error promotion model [38] proposes attention functions by increasing the precision (reliability) of prediction error signals [4, 37, 64]. This study lends support to the prediction error promotion model, since larger responses were elicited to unpredictable (for both the N170 and N2pc), and attended (only N170) stimuli, albeit not as an interaction. Under this model attention increases prediction errors by ensuring attention is directed to the most unpredictable (compared to predictable) of task relevant information to facilitate updating of priors, in order to reduce surprise (aka future occurrence of prediction error) when encountered in future. By doing so, attention (by increasing prediction error processing) operates by updating current statistical learning about goal-relevant information in the environment [38]. In other words, by updating our knowledge of the environment, we are strengthening associations between stimuli, making our future environment more “predictable”. This model is supported by previous studies involving multivariate pattern classifiers in fMRI [38], surprising visual stimuli using EEG [36], as well as in auditory stimuli using EEG scalp and source data [35]. By this view attention increases prediction error signals, and in doing so, enhances the distinction between expected and unexpected stimuli [38].

The current results suggest that whilst prediction and attention do modulate the sensitivity of ERP responses, it appears in this instance that endogenous cued stimuli separately influence attention and prediction in visual processing due to the lack of an interaction effect. The predictability effects on the N170 and N2pc are not contingent on the nature of endogenous attention required for the stimuli in this particular paradigm. These results have implications for future studies in using the N170 and N2pc for investigating the relationship between attention and predictive coding processing, as the N170 and N2pc appear not to reflect interactive effects for attention and prediction. The possibility remains that the present attention manipulation may not have been a sufficient manipulation as much as other visual studies using task relevant stimuli requiring a behavioural response to targets such as an appearance of faces or scenes [38] or a change in visual contrast [36]. Nevertheless, the motivation behind the cued stimuli in the present study was to manipulate attention in a more implicitly salient manner than in previous research.

## Conclusion

This study demonstrates prediction by prior context should be considered when investigating attention processes of the N170 and N2pc. The findings of this study reiterate the need to continue to control for prediction in attention components, as confirmed expectations demonstrated a dampening in N170 and N2pc responses evidenced by reduced amplitudes in predictable conditions. This research also highlights the value of prior context in influencing the amplitude of the N2pc component. Whilst the complicated nature of a conjunction of features and combination of attention and prediction factors encountered in daily life must be acknowledged, this research has demonstrated investigating visual perception using singleton feature stimuli assists by improving our knowledge of how attention and prediction direct processing of visual information, and why some stimuli may capture attention more so than others.

## Acknowledgments

We would like to thank all the research participants for generously donating their time to participate in this research, without whom this study would not be possible. We would also like to acknowledge the gentle encouragement of the Oily Rag Foundation (EN10007). K.S.B. was supported by an Australian Government Research Training Program Stipend.

## Supporting Information

**S1 Fig. Example of one Red Dot Sequence**. Example of one sequence for one of the red dot trials as the vigilance task. Red dots appeared randomly on any of the five steps in the sequence, as can be seen here on step three. Stimulus duration as indicated, 0ms intertrial interval, 500ms intersequence interval. All red dot trials were equally balanced across conditions and were discarded from EEG analyses on the basis of acting solely as a vigilance task to maintain gaze.

