## Supplementary figures and images for "Attention and prediction modulations in expected and unexpected visuospatial trajectories"

### Supplemental Fig 1

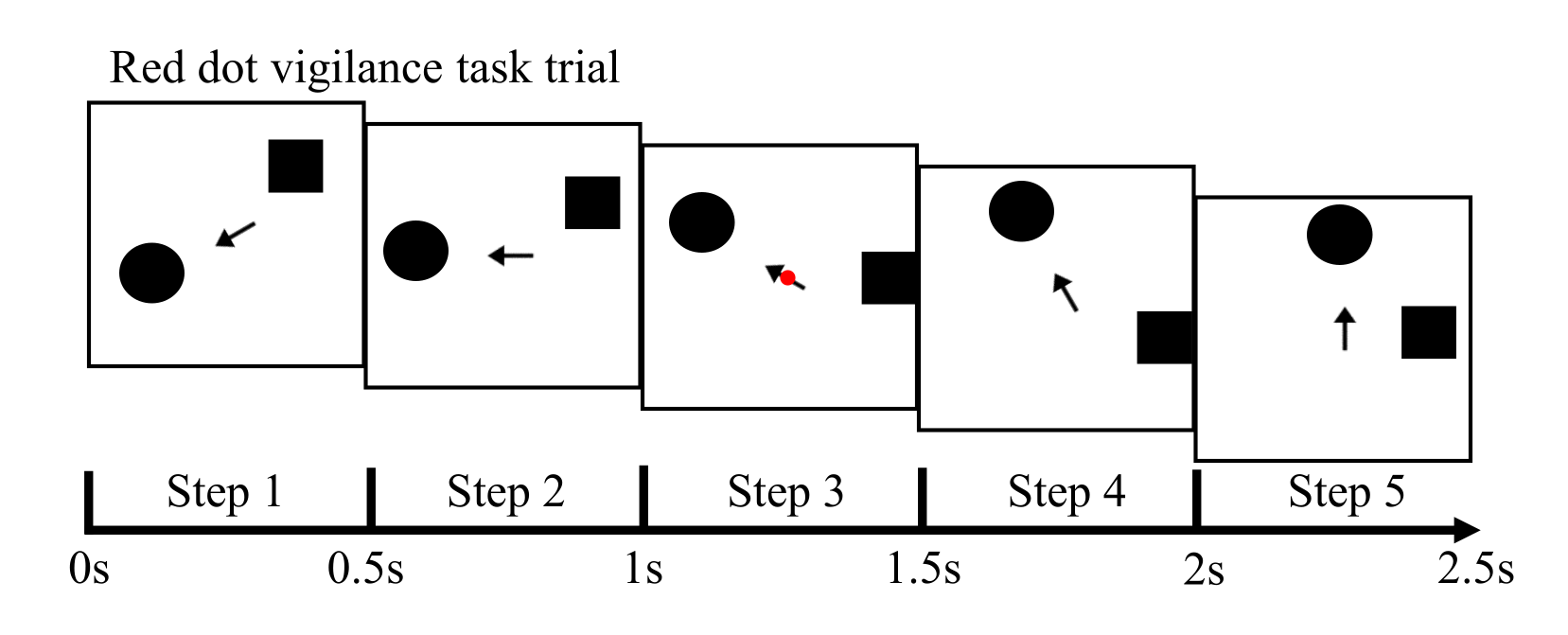
